# Anticipating on-target resistance to WRN inhibitors in microsatellite unstable cancers

**DOI:** 10.64898/2026.01.22.700152

**Authors:** Mark E. Orcholski, Nancy Laterreur, Wardah Masud, Shamika Shenoy, Vincent Chapdelaine-Trépanier, Julian Bowlan, Amisha Minju-OP, Júlia-Jié Cabré-Romans, Tanja Sack, Chris Fiore, Jordan T.F. Young, Alejandro Álvarez-Quilón, Raquel Cuella-Martin

**Affiliations:** Department of Human Genetics, McGill University, Montreal, QC, Canada; Victor Phillip Dahdaleh Institute of Genomic Medicine, McGill University, Montreal, QC, Canada; Repare Therapeutics Inc., Saint-Laurent, Quebec, Canada; DCx Biotherapeutics Inc., Saint-Laurent, Quebec, Canada; Servier Pharmaceuticals, Boston, MA, USA; AstraZeneca, Gaithersburg, MD, USA

**Author notes:** Equal contribution (Nancy Laterreur, Wardah Masud, Shamika Shenoy). Corresponding authors: Alejandro Álvarez-Quilón, Raquel Cuella-Martin.

**Keywords:** WRN, microsatellite instability (MSI), on-target drug resistance, synthetic lethality, HRO761, VVD-214, base editing, deep mutational scanning, CRISPR screens

## Abstract

Leveraging WRN helicase dependency in microsatellite instability (MSI) cancers offers a synthetic lethal (SL) therapeutic opportunity, with several WRN inhibitors in development. However, the hypermutator nature of MSI tumors creates strong evolutionary pressure for rapid resistance. Here, we apply a multimodal functional genomics framework integrating base editing screens and deep mutational scanning to map on-target resistance to two clinical WRN inhibitors, HRO761 and VVD-214. We identify discrete resistance hotspots within WRN and demonstrate that single-allele (heterozygous) mutations at the drug-binding site are sufficient to abrogate WRN inhibitor–induced cytotoxicity. Resistance profiles diverged between HRO761 and VVD-214, revealing mutations that impair one but preserve sensitivity to the other. Genome-wide CRISPR screens further identified non-homologous end joining (NHEJ) factors and the checkpoint phosphatase WIP1 as tractable synthetic vulnerabilities that potentiate WRN inhibition. Together, these findings establish a framework for resistance-aware deployment of WRN inhibitors through rational drug selection, therapeutic switching, and combination strategies.

**Statement of Significance:** Resistance to WRN inhibitors threatens the clinical durability of synthetic lethal therapies in microsatellite-instable cancers. Using multimodal functional genomics, we identify predictable, drug-specific on-target resistance mechanisms and reveal DNA-PK as a tractable combination partner. These findings provide a framework for resistance-aware deployment of WRN inhibitors to improve therapeutic durability.

## Introduction

Mismatch repair–deficient (MMRd) tumors, which account for a substantial fraction of colorectal (~15%), endometrial (~20–30%), and gastric (~10–20%) cancers, are molecularly characterized by microsatellite instability (MSI) (1). Loss of WRN is selectively lethal in MSI-high tumor cells but dispensable in MMR-proficient tissues, establishing WRN inhibition as a tractable synthetic lethal (SL) strategy with broad therapeutic potential (2–5). The biological basis of this dependency lies in the molecular consequences of MMR deficiency. Loss of the MMR genes *MLH1, MSH2, MSH6*, or *PMS2* drives hypermutation through the accumulation of insertion–deletion errors in microsatellite tracts, which are intrinsically prone to replication slippage due to their repetitive sequence composition (6). This process generates TA-rich sequences that form secondary DNA structures, causing replication fork stalling (7). MSI-high tumors, therefore, carry an intrinsic replication stress burden that normal cells lack. WRN helicase is uniquely required to unwind these structures and prevent catastrophic fork collapse, making its inhibition acutely toxic to MSI-high cells (7).

This dependency has generated substantial clinical interest, positioning WRN alongside other landmark SL paradigms—most notably PARP inhibition in BRCA1/2-mutant cancers. Multiple WRN inhibitors have been developed through independent pharmaceutical programs. Lead inhibitors target WRN allosterically, either through non-covalent (HRO761, Novartis; IDE275, Ideaya/GSK; NDI-219216, Nimbus Therapeutics) or covalent binding (VVD-214, Vividion Therapeutics; MOMA-341, Moma Therapeutics) (8–11). Clinical testing for HRO761 and VVD-214 began in 2023/2024 (NCT05838768, NCT06004245), with IDE275 (NCT06710847), NDI-219216 (NCT06898450), and MOMA-341 (NCT06974110) entering clinical trials in 2025.

This scenario highlights a pressing translational problem: The same hypermutator phenotype that generates WRN dependency in MSI-high cancers also provides the ideal evolutionary conditions for the rapid acquisition of drug resistance. Accordingly, xenograft studies of HRO761 already showed tumor rebound following an initial response (9). Yet, the on-target or off-target genetic alterations that confer resistance to WRN-targeted therapies—and can lead to clinical failure—remain incompletely defined. Other therapeutic approaches have previously shown that strong initial responses can mask rapid evolutionary escape in genetically unstable tumors, underscoring the need to anticipate therapeutic resistance to develop effective treatments (12). WRN inhibition triggers substantial replication stress and DNA damage, suggesting that MSI-high tumors may rely on compensatory repair and checkpoint pathways for survival. Identifying these pathways is critical to design rational combination strategies that enhance WRN-inhibitor efficacy and extend response durability.

Here, we employed a multimodal functional genomics strategy to systematically characterize resistance to WRN inhibition in MSI-high cancers. Using complementary base editing and deep mutational scanning approaches, we delineated the spectrum of WRN mutations capable of mediating on-target resistance and revealed that single heterozygous alterations at the drug-binding site are sufficient to confer resistance despite the presence of a wild-type allele. Importantly, resistance profiles differed across WRN inhibitors with distinct mechanisms of action, indicating that escape is not uniform and tumors may preserve sensitivity to alternative agents. Beyond on-target resistance, chemogenomic screening uncovered DNA repair and checkpoint pathways that modulate WRN dependency, highlighting opportunities for combination strategies to enhance efficacy and constrain resistance. Together, these findings underscore the substantial heterogeneity in WRN inhibitor sensitivity across MSI-high tumors and emphasize the importance of tumor genetic context and molecular profiling for informed therapeutic deployment.

## Results

### Cytosine and adenine base editing screens identify WRN mutations that cause resistance to the WRN inhibitor HRO761

To systematically identify WRN mutations that confer resistance to the WRN inhibitor HRO761, we performed cytosine and adenine base editing screens spanning the *WRN* coding region. The MSI-high colorectal cancer cell line HCT116 was engineered to stably express the cytosine base editor FNLS– NGG (HCT116–FNLS–NGG) or the adenine base editor TadA–NG (HCT116–TadA–NG). Polyclonal populations were then validated for their ability to introduce deleterious edits in common essential genes (**Fig. S1A-B**). Base editor-expressing cells were transduced with a single guide (sg)RNA library targeting every NGN protospacer adjacent motif (PAM) in *WRN* and were either left untreated or treated with HRO761 for 12 days to enrich for resistant variants. Screens were performed in biological triplicate, and log2 fold changes (LFCs) were calculated by comparing sgRNA abundance at the final versus initial timepoints (T18 *vs* T0) (**Fig. 1A, Table S1**). Quality control analyses confirmed high replicate concordance and strong recall in the FNLS–NGG screens (**Fig. S1C-D**). By contrast, TadA–NG screens showed reduced replicate correlation and recall, consistent with the lower editor efficiency and the limited deleteriousness of the iSilence positive control set, given that start codon removal does not reliably generate loss-of-function phenotypes (**Fig. S1A-D**).

**Figure 1.**
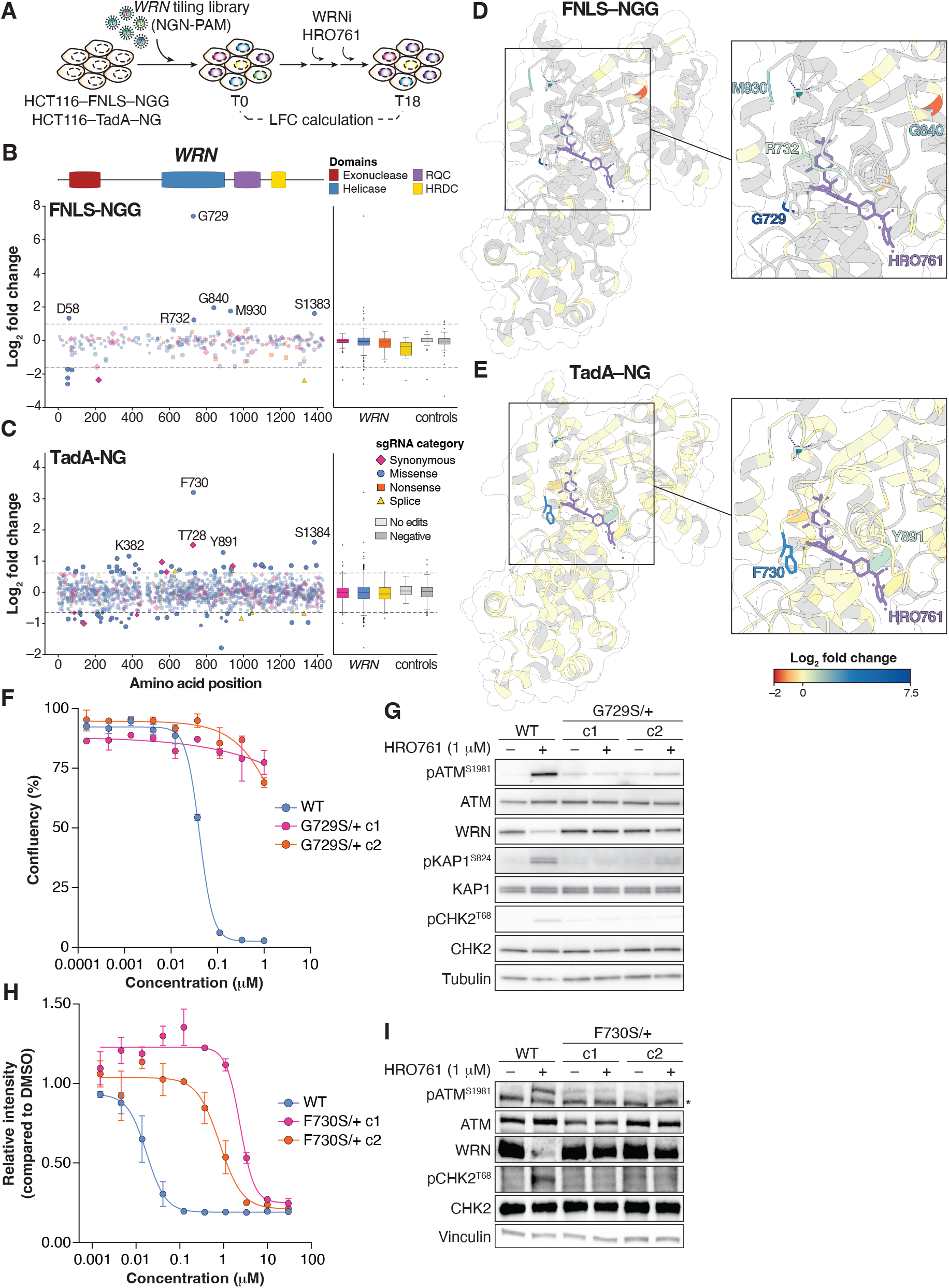
Base editing screens systematically identify mutations in WRN that cause resistance to HRO761. (**A**) Schematic of the base editing screens. (**B, C**) Base editing screen results in HCT116–FNLS–NGG (**B**) and HCT116–TadA– NG (**C**) cells treated with HRO761, mapped along the WRN protein as LFCs (T18 compared to T0). Dotted lines represent the threshold for biological relevance (1% of the negative control distribution), and symbol sizes represent statistical significance (large symbols indicate p-value < 0.01). sgRNAs are categorized by their predicted mutational outcomes, and the LFC distribution for each sgRNA category is depicted alongside those of empty-window (no edits) and designed negative controls (*AAVS1*-, *LacZ*-, and intron-targeting sgRNAs, as well as non-targeting sgRNAs). (**D, E**) Structure of HRO761 bound to WRN (PDB ID: 8PFO) (9) color-coded based on base editing screen phenotypes from (B, C). (**F**) Survival curves of HCT116 WT and two independent heterozygous clones carrying the G729S mutation (c1 and c2). Cells were exposed to HRO761 for 7 days, and confluency was measured using Incucyte. Mean ± s.e.m. N = 2. (**G**) Immunoblot of HCT116 WT and G729S/+ cells treated with HRO761 at 1 μM for 24 h. (**H**) Survival curve of HCT116 WT and two independent heterozygous clones carrying the F730S mutation (c1 and c2). Cells were exposed to HRO761 for 7 days, and confluency was measured by DAPI staining followed by total-intensity quantification. Mean ± s.e.m. compared to DMSO-treated control cells. N = 3. (**I**) Immunoblot of HCT116 WT and F730S/+ cells treated as in G.

SgRNAs predicted to introduce G729S and F730S mutations showed the strongest enrichment in the FNLS–NGG and TadA–NG screens, respectively (**Fig. 1B, C**). These residues lie within the HRO761 binding pocket on WRN, and their mutation is well positioned to disrupt the drug-target interaction (**Fig. 1D, E**). A nearby mutation predicted to generate R732Q also conferred modest resistance in the FNLS– NGG screen, consistent with local perturbation of the binding pocket (**Fig. 1B, D**). Beyond the immediate binding site, the screens revealed enrichment of sgRNAs introducing G840N, M930I/G931K, and Y891C/K892E substitutions within the helicase domain, suggesting potential allosteric mechanisms affecting inhibitor binding (**Fig. 1B-E**). Both screens also enriched sgRNAs inserting mutations in an evolutionarily conserved serine patch in the C-terminal region of WRN (S1383 and S1384), indicating that the responses to HRO761 involve regulatory elements outside the helicase domain (**Fig. 1B-C)**. Importantly, these mutations were enriched only under HRO761 treatment, with no observable phenotypes under untreated conditions (**Fig. S1E**).

### Heterozygous mutations within the WRN–HRO761 binding site cause resistance to WRN inhibition

To validate the screen results, we focused on the strong resistance-inducing mutations G729S and F730S, which were individually introduced into HCT116–FNLS–NGG and HCT116–TadA–NG cells, respectively. Interestingly, recovery of monoclonal populations yielded no homozygous clones for G729S; heterozygous G729S/+ clones were resistant to HRO761 compared to a WT control (**Fig. 1F, S1F**). Mechanistically, treatment with HRO761 activated ATM and downstream targets KAP1 and CHK2, and induced WRN degradation (**Fig. 1G**). These drug-induced phenotypes were completely abolished in the presence of the G729S mutation (**Fig. 1G**).

Inserting the F730S mutation proved inefficient in HCT116–TadA–NG cells with the sgRNA retrieved from the screen results (data not shown). To confirm that F730S was the causal driver of resistance, we therefore utilized the highly efficient ABE8e-SpG base editor together with three sgRNAs whose editing windows encompassed the F730 codon (**Fig. S1G**). Polyclonal populations edited at the target site were generated and treated with a high concentration of HRO761 (600 nM) for 8 days or left untreated. While synonymous mutations D731= and F730= were only modestly enriched during treatment, F730S was dramatically enriched in surviving cell populations (LFC > 3) relative to untreated controls, supporting its role as a strong resistance allele (**Fig. S1G**). Monoclonal populations were also recovered and genotyped (**Fig. S1H**). As with G729S, no homozygous F730S clones were recovered from either treatment-naïve or HRO761-treated polyclonal populations. Heterozygous F730S/+ clones were highly resistant to HRO761, with IC50 values approximately two orders of magnitude higher than WT cells (**Fig. 1H**). WRN F730S/+ cells were likewise resistant to HRO761-induced WRN degradation and failed to activate ATM-dependent signaling in the presence of the drug (**Fig. 1I**).

Collectively, these experiments demonstrate that base editing screens successfully mapped hotspots of WRN mutations conferring resistance to HRO761. Importantly, heterozygous G729S and F730S mutations were sufficient to induce resistance to WRN inhibitors, indicating that a single mutant allele may be enough to drive tumor resistance in patients.

### Deep mutational scanning of core WRN helicase and RQC regions reveals differential mutation patterns causing resistance to HRO761 and VVD-214

Data from base editing screens highlighted residues within the drug-binding pocket that are important for conferring drug resistance (**Fig. 1B-E**). Notably, the results also hinted at potential allosteric regulation within WRN’s helicase domain; however, the limited codon coverage and restricted mutational outcomes introduced with base editing may have prevented the identification of additional resistance-conferring sites. To generate a more comprehensive mutation–phenotype map of the helicase region surrounding the HRO761 binding pocket, we constructed a deep mutational scanning (DMS) library spanning WRN exons 19 (amino acids 697-758), 21-23 (amino acids 817-942) within the context of full-length WRN (**Table S2**). This library was ectopically expressed in WT HCT116 cells, and cells were treated at an LD50 concentration of HRO761 (40 nM) for 12 days (**Fig. 2A**). A DMSO-treated population served as a control. Variant-specific barcodes were sequenced at the final time point, and LFC values were calculated relative to the DMSO control. Similarly, the DMS library was probed against a second WRN inhibitor, VVD-214, at an LD50 (7 nM) for 12 days (**Fig. 2A**). Although HRO761 and VVD-214 bind the same pocket, we hypothesized that their distinct binding modalities would yield different resistance landscapes: tumors acquiring resistance to HRO761 might retain sensitivity to VVD-214, and *vice versa*. Consistent with the base editing results, mutations in G729 and F730 conferred resistance to HRO761 (**Fig. 2B**). A second hotspot centered on V922 also emerged, identifying additional residues essential for preserving the integrity of the HRO761-binding pocket (**Fig. 2D**). Mutations conferring resistance to VVD-214 likewise localized to the drug-binding pocket, particularly at C727, G729, and to a lesser extent, F730 (**Fig. 2C,E**). Distinct from HRO761, the VVD-214–WRN interaction also involves E846, whose mutation produced a strong resistance phenotype (**Fig. 2D,E**). Additional resistance-causing mutations were detected in residues adjacent to or even distal from the binding pocket, including E851 and D832, respectively (**Fig. 2C,E**).

**Figure 2.**
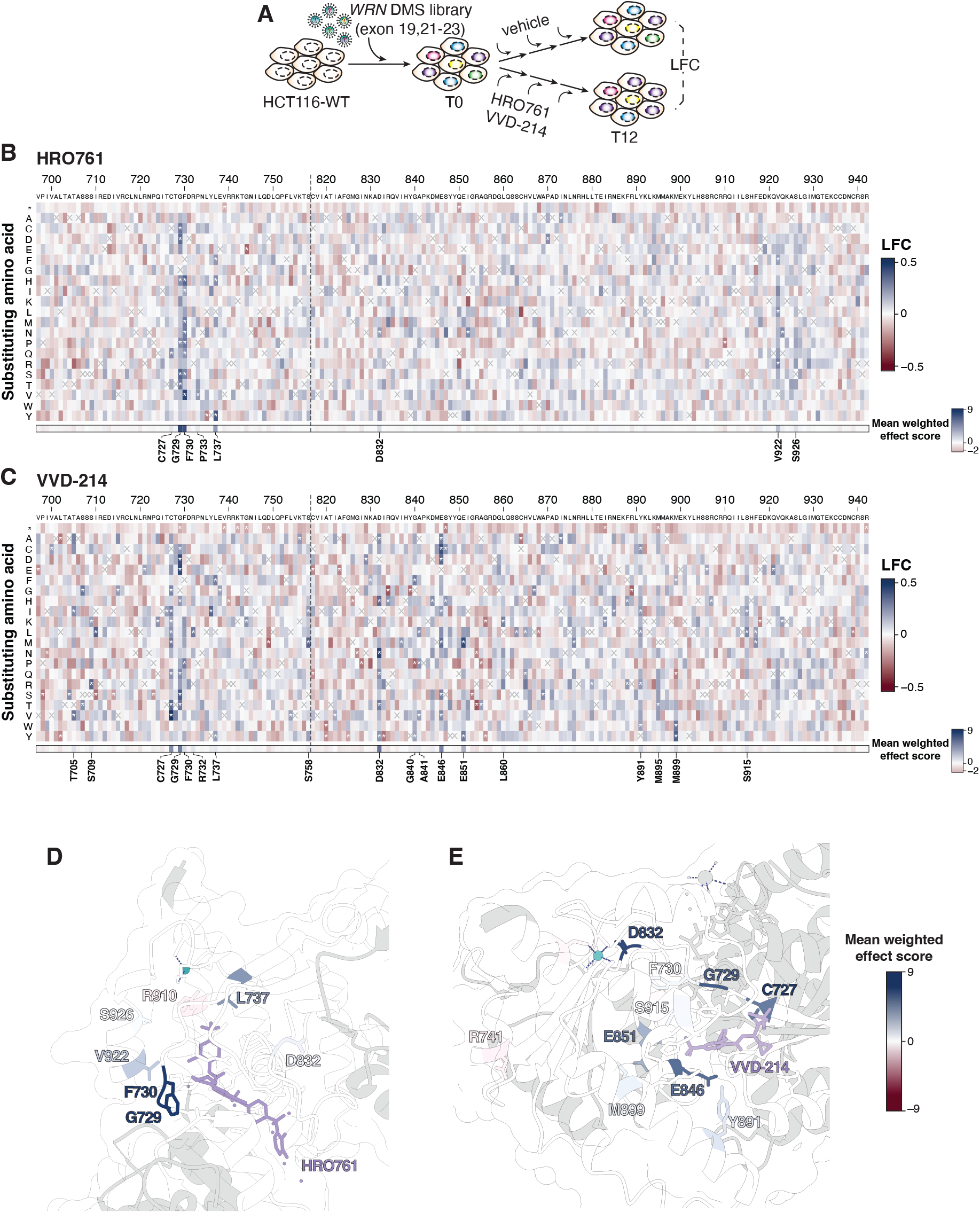
Deep mutational scanning reveals different patterns of on-target resistance to HRO761 and VVD-214. (**A**) Schematic of the DMS experiment. (**B, C**) Heatmap representation of DMS results across the WRN helicase region encoded by exons 19, 21-23 following treatment with HRO761 (**B**) or VVD-214 (**C**). Dotted lines indicate a discontinuous sequence. LFCs were calculated by comparing inhibitor-treated to DMSO-treated cells at the endpoint. Weighted effect scores per mutation are computed as the product of LFC and the negative log10 of the p-value. The per-residue mean of the weighted effect scores was then standardized, such that each reported value reflects the number of standard deviations by which that mean differs from the overall mean. Asterisks (*) indicate statistically significant changes, and the (X) marks the WT amino acid. (**D, E**) Structures of WRN bound to HRO761 (PDB ID: 8PFO) (9) (D) or VVD-214 (PDB ID: 7GQU) (8) (E), colour-coded by per-residue mean weighted effect score (standardized). Amino acids with the highest absolute effect scores are highlighted.

We next sought to validate the similarities and differences between the resistant mutation profiles for HRO761 and VVD-214, and therefore selected four representative mutations: G729S, which conferred resistance to both drugs; C727V and E846S, which specifically conferred resistance to VVD-214; and F730H, which appeared to confer resistance only to HRO761 based on the DMS data (**Fig. S2B**). WRN WT and mutant open-reading frames were expressed in HCT116 cells (**Fig. S2C**), and dose-response analyses were performed. Expression of WRN G729S led to a ~2-fold increase in IC50 for HRO761 compared to WT cells; resistance to VVD-214 was even more pronounced, showing a ~10-fold increase in IC50 compared to WT cells (**Fig. 3A**). Molecularly, WRN G729S was resistant to degradation after treatment with HRO761, and ATM activation and H2AX phosphorylation (γH2AX) were nearly abolished (**Fig. 3B, C**). Despite the observed resistance upon VVD-214 treatment, ATM activation was mildly diminished, and H2AX was significantly phosphorylated in cells carrying WRN G729S (**Fig. 3C**), indicating that G729 may be more critical for HRO761 than VVD-214 binding.

**Figure 3.**
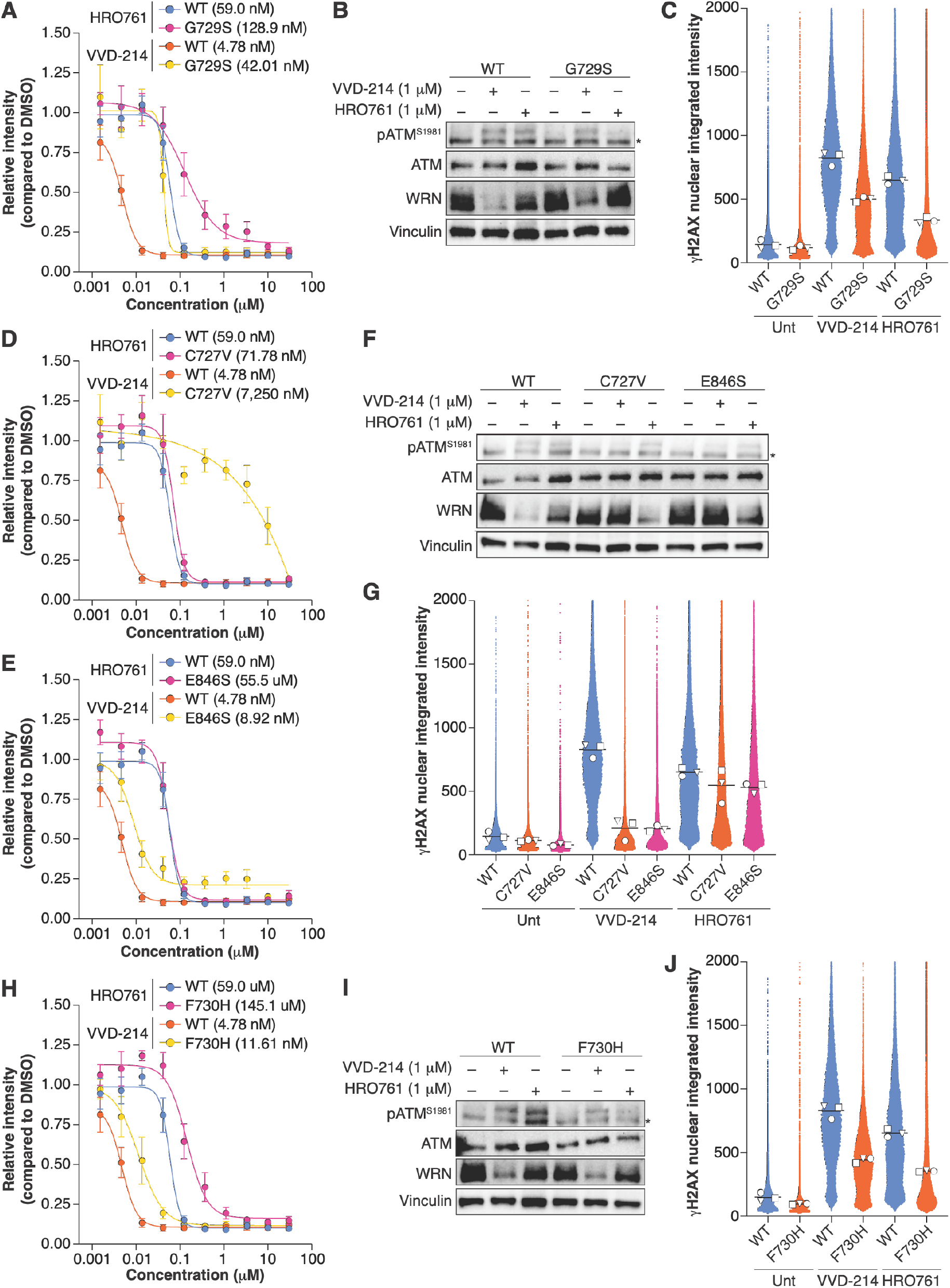
Mutations conferring resistance to VVD-214 remain sensitive to HRO761. (**A, D, E, H**) Survival curves of HCT116 cells overexpressing WT or G729S (A), C727V (D), E846S (E), or F730H (H) WRN. Cells were exposed to HRO761 or VVD-214 for 7 days, and confluency was measured by DAPI intensity. Mean ± s.e.m. compared to DMSO-treated control cells. N = 3-5. (**B, F, I**) Immunoblot of HCT116 WT cells overexpressing WT or G729S (B), C727V (F), E846S (F), or F730H (I) WRN, untreated or treated with VVD-214, HRO761 at 1 μM for 24 h. (**C, G, J**) Quantification of the integrated intensity per nucleus of γH2AX immunostaining in basal conditions or upon treatment with HRO761 (10 μM) and VVD-214 (1 μM) in HCT116 cells overexpressing WT or G729S (C), C727V (G), E846S (G), or F730H (J) WRN. Mean (N = 3), with a minimum of 10,000 cells per biological replicate, is depicted.

Consistent with the role of C727 in covalent binding of VVD-214 (8), introduction of the C727V mutation produced a >100-fold increase in the IC50 for VVD-214 compared to WT. In contrast, no difference was observed in the presence of HRO761 (**Fig. 3D**). Similarly, the E846S mutation conferred specific resistance to VVD-214, albeit to a lesser extent (~2-fold, **Fig. 3E**). Interestingly, both mutations conferred similar phenotypic results at a molecular level: no VVD-214-dependent degradation was triggered, and ATM activation and H2AX phosphorylation appeared nearly abolished in cells carrying C727V and E846S WRN (**Fig. 3F-G**). HRO761 treatment in these cells was able to include ATM activation signalling and H2AX phosphorylation (**Fig. 3F-G**).

DMS data suggested that F730H would confer specific resistance to HRO761. Drug-response analyses showed a comparable increase in resistance to both HRO761 and VVD-214 (~2.5-fold relative to WT cells) (**Fig. 3H**). However, molecular analyses revealed WT-like degradation of WRN F730H upon treatment with VVD-214 and partially preserved ATM-dependent signaling, highlighting F730H as a hotspot for HRO761 resistance (**Fig. 3I-J**).

Collectively, these results highlight differences in the resistance profiles for HRO761 and VVD-214, suggesting that tumors acquiring resistance to one inhibitor through WRN mutation may remain sensitive to the other.

### Inhibition of DNA-PK and WIP1 synergizes with WRN inhibition to treat MSI-high cells

In addition to mutations in WRN itself, other cellular alterations may also influence tumor response to WRN inhibitors. We stratified cell lines in the CRISPR Dependency Map by microsatellite status and examined the relationship between WRN dependency and sensitivity to HRO761. As expected, MSI-high cell lines displayed greater dependency on WRN than microsatellite stable (MSS) cells and exhibited lower AUC values, indicating higher sensitivity to HRO761 (**Fig. 4A**). However, a wide range of sensitivities to HRO761 was observed within the MSI-high group. This heterogeneity was validated in a panel of six MSI-high and six MSS/MSI-low cell lines (**Fig. 4B**). While HCT116, SW48, and LS411N were highly sensitive to HRO761, LoVo and RKO cells showed only intermediate sensitivity (**Fig. 4B**). Notably, direct comparison between a sensitive MSI-high model (HCT116) and a less sensitive model (RKO) revealed a >5-fold difference in their HRO761 IC50 values (**Fig. S3A**).

**Figure 4.**
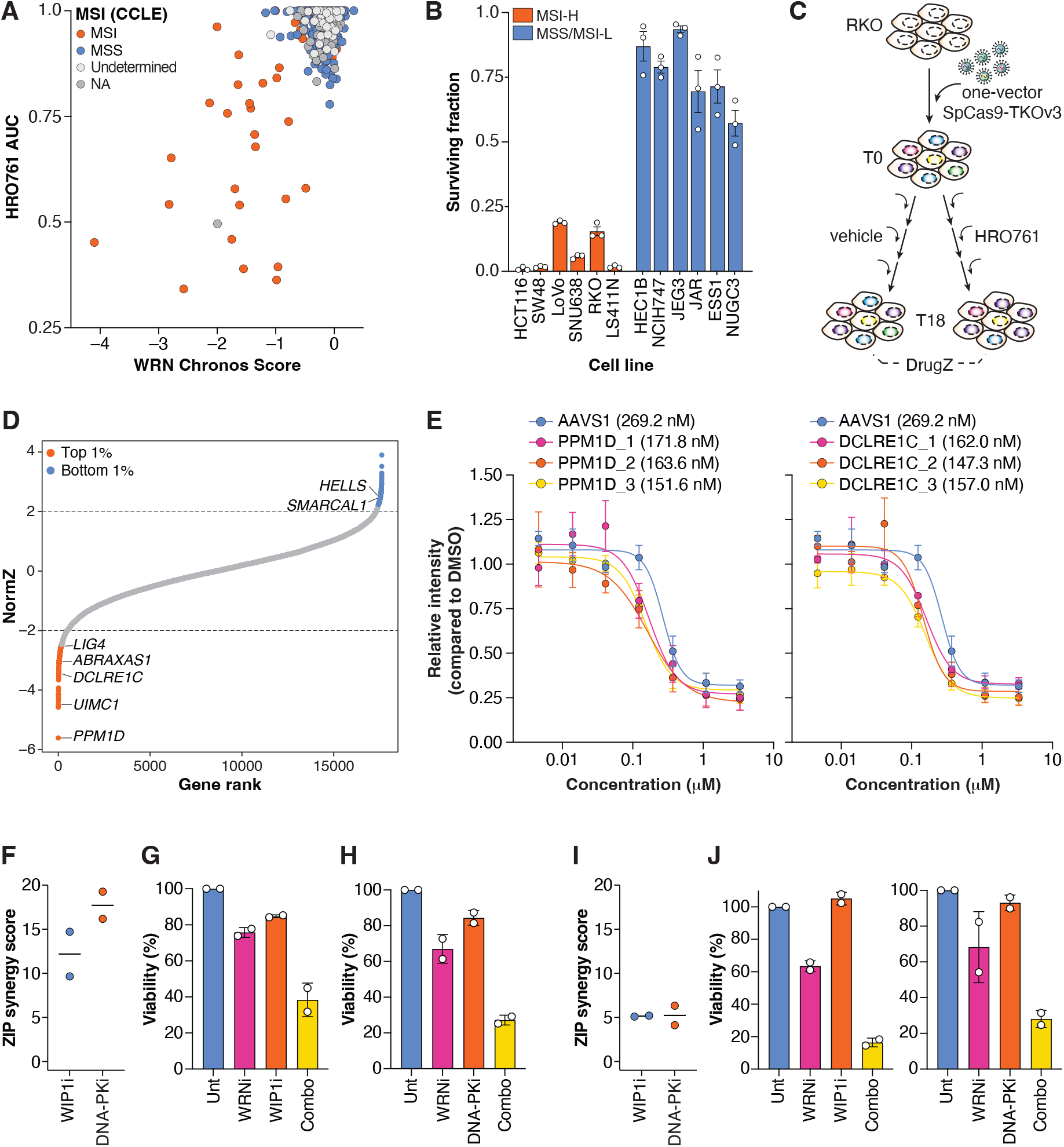
WIP1 and DNA-PK inhibitors synergize with HRO761 to induce cell death in MSI-high cell lines. (**A**) Analysis of cell-line dependency on WRN and susceptibility to HRO761, stratified by MSI status; MSS: microsatellite stable; NA: non-applicable (**Table S3**). (**B**) Clonogenic assays on a representative panel of MSI-high and MSS/MSI-low cell lines in response to HRO761 (1 μM). The X axis represents the surviving fraction compared to the DSMO control counts. (**C**) Schematic of the genome-wide synthetic lethality screen design. (**D**) Representation of NormZ values per gene generated using DrugZ, comparing DMSO-treated versus HRO761-treated RKO cell populations. (**E**) Survival curves of RKO cells transduced with sgRNAs targeting *PPM1D* (left) and *DCLRE1C* (right) using a one-vector system. Six days post-infection, cells were exposed to HRO761 for 7 days, and confluency was measured by total DAPI intensity. Mean ± s.e.m. compared to DMSO-treated control cells. N = 4; except DCLRE1C_1, N = 2. **(F)** ZIP synergy scores derived for the combination of HRO761 with either GSK-2830371 (WIP1i) or AZD-7648 (DNA-PKi) in RKO using the matrix datasets in Fig. S3D. Mean for N = 2 (**G**) Survival analysis of RKO cells treated with either HRO761 (WRNi) at 179 nM, GSK-2830371 (WIP1i) at 5 μM or both (combo) for 7 days. Confluency was measured with Incucyte. Relative survival is calculated with respect to a DMSO-treated control. Mean ± SD for N = 2. (**H**) Survival analysis of RKO cells treated with either HRO761 at 89 nM, AZD-7648 (DNA-PKi) at 1.78 μM or both for 7 days, as in (**G**). Mean ± SD for N = 2. (**I**) ZIP synergy scores derived for the combination of HRO761 with either GSK-2830371 (WIP1i) or AZD-7648 (DNA-PKi) in HCT116 cells. Mean for N = 2 (**J**) Survival analysis of HCT116 cells treated with either HRO761 at 70 nM, GSK-2830371 at 1.6 μM or both; HRO761 at 45 nM, AZD-7648 (DNA-PKi) at 6.45 μM or both for 7 days. Mean ± SD for N = 2.

We hypothesized that, as in cell lines, the genetic makeup of MSI-high tumors may further modulate clinical responses to WRN inhibitors. Accordingly, specific genetic vulnerabilities might synergize with WRN inhibition and highlight combination therapy strategies capable of broadening WRN inhibitor efficacy across MSI-high tumors. Conversely, genetic alterations causing WRN inhibitor resistance can highlight specific off-target resistance mechanisms that may pre-exist or arise in MSI-high tumors. To test this, RKO cells were transduced with the genome-wide, all-in-one CRISPR-knockout library TKOv3. Cells were left untreated or treated with HRO761 at an LD20 for 18 days to identify both potential resistance and sensitivity-conferring genes. HRO761-specific genetic interactions were identified by comparing the treated versus untreated endpoints using DrugZ (**Fig. 4C**). Consistent with previous reports (13), the screen did not reveal strong genetic interactions driving off-target therapeutic resistance, potentially due to the HRO761 dose used or masked by pre-existing alterations in RKO cells that already confer a degree of intrinsic resistance. Notably, among the top hits we found *HELLS* and *SMARCAL1*, recently identified as modulators of WRN inhibitor activity (14). Among the top sensitizing hits were the phosphatases PPM1D (WIP1) and PPM1G, both involved in p53 regulation, as well as members of the BRCA1-A complex (*UIMC1*/RAP80, *ABRAXAS1*) and components of the non-homologous end-joining pathway (*DCLRE1C*/Artemis, *LIG4*) (**Fig. 4D**). Because WIP1 and NHEJ inhibitors—particularly DNA-PK inhibitors—have been explored as anti-tumorigenic agents (15,16), we focused on validating *PPM1D, DCLRE1C*, and *LIG4* using survival-based assays. Polyclonal knockout populations of *PPM1D, DCLRE1C* and *LIG4*-targeted cells using independent sgRNAs exhibited ~1.5 to 1.8-fold increased sensitivity to HRO761 compared to an *AAVS1*-targeted control (**Fig. 4E, S3B**). All sgRNAs edited their corresponding locus at high efficiency (**Fig. S3C**).

To determine whether pharmacological inhibition of PPM1D recapitulated this phenotype, we employed the WIP1 inhibitor GSK-2830371 (17) and probed a combinatorial range of concentrations. Synergy was then computed using the zero interaction potency (ZIP) reference model (18), resulting in an average score of 12.19 on RKO cells, consistent with a synergistic effect (**Fig. 4F**). While treatment with GSK-2830371 achieved optimal synergy with HRO761 when both were used at LD20 concentrations (**Fig. 4G**), synergies were observed across a range of concentrations (**Fig. S3D**). We next sought to achieve selective NHEJ inhibition by targeting DNA-PK with the potent DNA-PK inhibitor AZD-7648 (19). Combining DNA-PK and WRN inhibition showed a strong synergistic effect, with an average ZIP synergy score of 17.72 (**Fig. 4F, 3E**). Treatment with AZD-7648 at an ~LD20 concentration optimally synergized with HRO761 at an LD35 concentration, an effect also observed across a range of concentrations (**Fig. 4H, S3E**).

Notably, a modest synergy was also observed in HCT116 cells for both drugs tested (**Fig. 4I, J**), indicating that the combination could be applied across MSI tumors to potentiate the action of WRN inhibition.

## Discussion

Rather than asking whether resistance to WRN inhibition can occur, this study defines how accessible resistance routes are, which molecular mechanisms can mediate escape, and how such routes might be anticipated and constrained. A central challenge in targeting SL dependencies in MSI-high cancers is the paradox created by their hypermutator state: the same genomic instability that renders these tumors exquisitely sensitive to pathway inhibition also creates a permissive landscape for the emergence of resistance. Using complementary base editing and saturation mutagenesis approaches, we demonstrate that resistance to WRN inhibition can be mediated by on-target mutations within WRN itself, and that single-allele substitutions at key residues are sufficient to abrogate drug sensitivity. These findings indicate that the acquisition of a single drug-resistant allele substantially lowers the genetic barrier to escape treatment. Importantly, the resistance landscape is not uniform across inhibitors, revealing drug-specific rather than class-wide mutational escape routes.

To define these resistance routes, we systematically mapped resistance-conferring WRN mutations for two lead clinical inhibitors, HRO761 and VVD-214, using cytosine and adenine base editing screens in parallel with high-resolution DMS. Together, these complementary approaches delineate the spectrum of WRN substitutions capable of mediating resistance and reveal mechanistic heterogeneity in inhibitor escape. HRO761 inhibits WRN by non-covalently binding an allosteric site between the D1 and D2 helicase domains (9). Consistent with this mechanism, the strongest resistance mutations identified by base editing, G729S and F730S, clustered within the HRO761 binding interface. Functional validation showed that introduction of either mutation, even in heterozygosity, was sufficient to abrogate HRO761-driven cytotoxicity, while preserving cell viability, consistent with retained WRN function. Notably, despite efficient editing and drug selection, homozygous mutant clones were not recovered, suggesting that complete loss of wild-type WRN may be poorly tolerated under these conditions. These observations support a model in which WRN is haplosufficient for its essential cellular functions, such that acquisition of a single drug-resistant allele provides an efficient route to escape while maintaining helicase activity, thereby substantially lowering the genetic barrier to resistance in MSI-high tumors.

Mutations outside the HRO761–WRN binding interface, including G840N, M930I/G931K, and mutations within a conserved C-terminal serine patch (S1383G and S1384G), also conferred resistance in base editing screens. Together with pocket-disrupting substitutions, these findings indicate mechanistic heterogeneity in WRN inhibitor resistance, arising both from direct disruption of drug-target contacts and from distal mutations that likely alter WRN conformation or modulate accessibility of the binding site. Inhibition of WRN has been shown to trigger protein degradation, as inactive WRN is recognized as defective by the cellular quality-control machinery (8–9), thereby amplifying functional loss and enhancing selective toxicity in MSI cells. Whether resistance-conferring mutations outside the binding pocket influence WRN degradation upon inhibition remains an open mechanistic question.

Base editing approaches are inherently constrained in mutational scope, as they only introduce specific nucleotide transitions and depend on compatible PAM sequences (20). To achieve more comprehensive coverage of WRN resistance space, we performed DMS across the WRN helicase domain. While G729 and F730 again emerged as dominant resistance nodes for both HRO761 and VVD-214, DMS revealed a broader spectrum of residues, including V922, E846, and C727, whose substitution differentially impacted sensitivity to the two inhibitors. Notably, VVD-214 inhibits WRN through covalent modification of Cys727 within the same allosteric pocket targeted by HRO761, whereas HRO761 binds non-covalently. Consistent with these distinct binding modalities, mutations at C727 and E846 selectively conferred resistance to VVD-214, while preserving sensitivity to HRO761. This divergence in resistance profiles indicates that tumors escaping one WRN inhibitor may remain vulnerable to alternative inhibitors with distinct mechanisms of action, highlighting the potential for therapeutic sequencing or switching to circumvent acquired on-target resistance. Together, these complementary mutational approaches define the breadth of WRN resistance mutations, expose mechanistic heterogeneity between clinical WRN inhibitors, and provide a framework for rational deployment of WRN-targeted therapies.

Despite promising preclinical activity (8–11), ongoing clinical proof-of-concept studies are showing limited patient responses (21). The durability of WRN inhibitor responses as monotherapy also remains unknown, highlighting the importance of understanding and anticipating resistance mechanisms. Although WRN dependency is a defining feature of MSI-high tumors, dependency strength varies across MSI-high colorectal cancer cell lines, supporting the rationale for combination strategies to enhance efficacy and durability of response. Leveraging genome-wide CRISPR-KO chemical genetic screens, we identified *PPM1D* (WIP1) and multiple components of the non-homologous end joining (NHEJ) pathway, including *DCLRE1C* and *LIG4*, as sensitizers to WRN inhibition. Functional validation confirmed that pharmacological inhibition of WIP1 and DNA-PK, using GSK-2830371 and AZD-7648, respectively, potentiates the effect of HRO761 in suppressing tumor cell viability. Given the wild-type *TP53* status of both RKO and HCT116 cells, the response to WIP1 inhibitors is likely due to sustained p53 activity and induction of p53-dependent cell death (22), limiting its applicability to *TP53*-mutant tumors. Notably, the drug combination benefit was most pronounced with DNA-PK inhibition, consistent with a model in which WRN inhibition exacerbates replication-associated DNA damage that becomes critically dependent on NHEJ-mediated repair. Together, these findings support rational combination strategies that target collateral DNA repair vulnerabilities unmasked by WRN inhibition. Combination therapy could be positioned to constrain resistance rather than to simply amplify cytotoxicity in MSI-high tumors.

Finally, MSI-high tumors are highly immunogenic due to their elevated neoantigen burden, and immune checkpoint blockade—particularly PD-1/PD-L1 and CTLA-4 inhibitors—has transformed the clinical management of MSI-high colorectal cancers by delivering durable responses and improved survival (23–24). While immunotherapy has become the standard of care in this setting, not all patients achieve sustained benefit, highlighting the need for complementary therapeutic strategies. Although WRN inhibition represents a promising addition to the MSI treatment landscape (25), our findings suggest it cannot be optimally deployed as a stand-alone therapy and instead requires resistance-aware development strategies incorporating early mutation monitoring, rational drug switching, and pathway-informed combination regimens. Future efforts should focus on clinical detection of on-target WRN resistance mutations and development of next-generation WRN inhibitors with complementary resistance profiles. Additional efforts should also be made to evaluate WRN-based combination regimens, including both chemotherapy and immunotherapy, in prospective clinical trials aimed at maximizing response durability for patients with MSI-high tumors.

## Methods

### Cell lines and cell culture

HEK293T, HCT116, and SW48 cell lines were cultured in high-glucose DMEM supplemented with 2 mM L-glutamine (Gibco, Thermo Fisher Scientific), 10% (v/v) fetal bovine serum (Gibco, Thermo Fisher Scientific), and 100 U/mL penicillin plus 100 μg/mL streptomycin (Thermo Fisher Scientific). LOVO, SNU638, RKO, LS411N, NCIH747, ESS1 and NUGC3 cell lines were cultured in RMPI supplemented as above. HEC1B cells were cultured in EMEM supplemented as above. All cells were maintained in a humidified incubator at 37 °C and 5% CO_2_.

HCT116–FNLS–NGG and HCT116–TadA–NG were generated by lentiviral transduction. Briefly, viral supernatants (VectorBuilder) were applied directly to HCT116 cells with 4 μg/ml polybrene for 16 h. Cells were then treated with blasticidin at 10 μg/mL for 5 days. FNLS–NGG and TadA–NG activity was determined by transfecting cells with synthetic sgRNAs predicted to insert known deleterious or neutral mutations using Lipofectamine CRISPRMAX (Thermo Fisher Scientific) according to the manufacturer’s instructions. Cell confluency was measured using Incucyte.

The HCT116–ABE8e–SpG cell line was generated by lentiviral transduction. Briefly, 5 × 10^6^ HEK293T cells were seeded in 10-cm dishes in antibiotic-free DMEM. Sixteen hours later, cells were co-transfected with 8 μg ABE8e–SpG–P2A–BlastR plasmid together with third-generation packaging plasmids gag-pol, VSV-G, rev, and tat (0.4 μg, 0.8 μg, 0.4 μg, and 0.4 μg) using TransIT-293 (Mirus Bio) according to the manufacturer’s instructions. Viral supernatants were collected at 48 and 72 h post-transfection, filtered through a 0.45 μm filter, and applied directly to HCT116 cells in a 1:1 virus/medium mixture growing with 4 μg/mL polybrene for 16 h. After two rounds of transduction, cells were treated with blasticidin at 10 μg/mL for 5 days. HCT116–ABE8e–SpG cells were subsequently maintained at 5 μg/mL blasticidin. ABE8e-SpG expression was confirmed by Cas9 immunoblotting.

Heterozygous HCT116–FNLS–NGG WRN G729S mutants were generated by transfecting a synthetic sgRNA targeting G729S (**Table S5**) using Lipofectamine CRISPRMAX (Thermo Fisher Scientific). Cells were seeded at low density, and monoclonal populations were recovered and validated by targeted PCR amplification and Sanger sequencing. Heterozygous HCT116–ABE8e–SpG F730S mutants were generated by lentiviral delivery of pLenti–Guide–Purov2 containing F730S–sgRNAs 1–3 (**Table S5**) using a single round of infection at a multiplicity of infection (MOI) > 1. Cells were selected with 2 μg/mL puromycin for 48 h, then maintained at 1 μg/mL puromycin for an additional four days. A T0 polyclonal population was collected at this stage. For enriched selection, cells were seeded at low density to recover monoclonal populations (treatment-naïve) or seeded 2 × 10^5^ cells per well in 6-well plates and treated with 600 nM HRO761 for 8 days (T8) prior to low-density plating. Polyclonal and monoclonal populations were characterized by targeted PCR amplification and Sanger sequencing.

HCT116 cells expressing WT or mutant WRN open-reading frames were generated by lentiviral transduction as described above. Viral supernatants were applied for a single round of infection, followed by selection with 2 μg/mL puromycin for 48 h, then maintenance at 1 μg/mL puromycin. WRN expression was determined by immunoblot.

### Plasmids

The FNLS–NGG–P2A–BlastR construct used in this study was built by VectorBuilder using the FNLS-NGG sequence from the FNLS–NGG–P2A–PuroR (26) expressed under the EF1a promoter. Similarly, the TadA-NG–P2A-BlastR was built using the ABE8e-NG sequence driven by the EF1a promoter. The ABE8e–SpG–P2A–BlastR was generated from pRDA_479 (a gift from John Doench & David Root, Addgene #179099, 27) and FNLS–NGG–P2A–BlastR (28) using NEBuilder HiFi DNA Assembly (New England Biolabs).

To generate the WRN F730S mutant, sgRNAs were cloned into a modified version of pLenti–Guide– Puro (a gift from Feng Zhang, Addgene #52963) (29) carrying the sgRNA scaffold from LRT2B (a gift from Lukas Dow, Addgene #110854) (26) to increase editing efficiency (hereafter referred to as pLenti– Guide–Purov2). For characterization of individual sgRNA-induced knockouts, sgRNAs (**Table S5**) were cloned into the pLenti–CRISPRv2 vector (a gift from Feng Zhang, Addgene #52961) (29). Briefly, complementary oligonucleotides containing BsmBI-compatible overhangs were synthesized (Integrated DNA Technologies), annealed, and ligated into BsmBI-digested sgRNA-expression plasmids using T4 DNA Ligase (New England Biolabs). sgRNA insertion was verified by Sanger sequencing (Genome Québec).

WT and mutant WRN open reading frames were synthesized and cloned into pLV[Exp]-Puro-CMV (VectorBuilder).

Plasmid DNA was subsequently prepared using the ZymoPure II Midiprep Kit (Zymo Research) for downstream transfections. All plasmids were verified by full plasmid sequencing (Plasmidsaurus Inc.).

### Drugs

HRO761 and VVD-214 were purchased from Ambeed, Inc., resuspended in DMSO at a concentration of 10 mM and stored at −80 °C. GSK-2830371 and AZD-7648 were acquired from MedChemExpress.

### Base Editing Screens

#### sgRNA library design

The sgRNA library consisted of WRN targeting guides, negative control guides, and positive control guides. To select sgRNAs targeting the WRN gene, we identified all possible guides in exonic regions and up to ± 40 nt spanning introns with an NGN PAM sequence (n = 2645). The negative control guides consisted of guides targeting introns of essential genes (n = 18), guides not targeting any gene (n = 21), and guides expected to lead to silent changes with a base editor to essential genes (n = 6). The positive control guides consisted of guides leading to a premature stop codon in essential genes (iSTOP (30), n=70 for NGG PAMs, n=29 for other NGN PAMs, and n=26 for other PAMs), guides leading to the removal of a start codon in essential genes (iSilence (31), n = 11 for NGG PAMs, n = 11 for other NGN PAMs). There were also 26 guides leading to premature stop codons and 3 guides leading to the removal of start codons in three genes that are expected to lead to resistance to WRN inhibitors when knocked out. Each guide was designed as a 19-bp sequence and preceded by a G to ensure transcription under the U6 promoter. The sgRNA library was cloned by VectorBuilder into an sgRNA expression vector with a PGK-Puromycin resistance marker.

#### Variant annotation

Mutational outcomes inserted by FNLS-NGG and TadA-NG were predicted using custom R scripts based on the crisprVerse set of packages (32). addEditedAlleles() from the crisprDesign package was used to determine all possible edited alleles and their relative probabilities. For this purpose, the editing weights of the TadA enzyme were set for ‘A2G’ as [0.05, 0.3, 0.8, 1, 1, 0.82, 0.55, 0.3, 0.2] for positions –19 to –11 relative to the PAM sequence. For the FNLS enzyme, the editing weights were set for ‘C2T’ as [0.05, 0.20, 0.44, 0.91, 1, 0.78, 0.41, 0.14, 0.07] for positions –19 to –11 relative to the PAM sequence. For each guide, the edited alleles were summarized by determining the most likely edit type (missense, nonsense, splice, no change) and the single most likely edit based on the resulting amino acid sequence of the edited allele. For edited alleles resulting in the same amino acid sequence, their scores were combined. Splice events were determined based on the canonical AG/GT splice site only. A minimum edited score of 0.1 (out of 1) was required to consider an edited allele; all other edited alleles were considered as non-occurring.

#### Base editing screen experiments

Base editing tiling screens were performed using protocols adapted from published literature (28,33,34). Briefly, HCT116 cells expressing FNLS–NGG or TadA–NG were transduced with lentivirus encoding a tiling sgRNA library for the *WRN* gene (all exons and exon-intron junctions) at an MOI of ~0.3. The screen was conducted in technical triplicates, and library coverage of >1,000 cells per sgRNA was maintained. Puromycin-containing medium (2 µg/mL) was added 2 days after infection to select for transductants. Selection was continued until 96 h after infection, which was considered the initial time point (T0). HRO761 was added to the cells at day 6 (T6) and day 10 (T10) at 60 nM and 10 nM at day 14 (T14), and the screen was terminated at day 10 (T18). Genomic DNA was isolated using the QIAamp Blood Maxi Kit (Qiagen), and sgRNA sequences were amplified by PCR using NEBNext Ultra II Q5 Master Mix (New England Biolabs). The i5 and i7 multiplexing barcodes were added in a second round of PCR, and final gel-purified products were sequenced on an Illumina NextSeq500 system at the Lunenfeld-Tanenbaum Research Institute (LTRI-NBCC).

#### Data analyses and representation

The number of reads per sgRNA was computed using the MAGeCK count command (35). SgRNAs with read counts below 200 in any condition were eliminated to prevent artifactual LFC values. Comparisons of T0 *vs* T18 were performed using paired MAGeCK robust rank aggregation (RRA) (e.g., mageck test -k Countfile.txt -t T18_rep1, T18_rep2, T18_rep3 -c T0_rep1, T0_rep2, T0_rep3 --paired -n T18 --adjust-method fdr). Biological significance thresholds were set at the top and bottom 1% values of ranked LFC for the negative controls. For statistical significance, the threshold was set to a one-sided p value < 0.01. SgRNAs meeting both criteria were considered hits.

Data analyses and representations were performed in Python (version 3.12.2) using matplotlib (3.10.3), seaborn (0.13.2), SciPy (1.15.3), and scikit-learn (1.7.0) packages. Repeat-to-repeat Pearson’s r correlation values were computed using per-repeat LFCs, and receiver operator characteristic–area under the curve (ROC–AUC) analyses were computed using negative MaGeCK rank values. Lollipop plots were generated using exclusively sgRNAs mapping and annotated to the *WRN* NM_000553.6 transcript and targeting sites matching the corresponding base editor PAM.

LFC data were mapped to the WRN structure bound to HRO761 (PDB ID: 8PFO, 9) using UCSF ChimeraX version 1.10rc202506130232 (36). Briefly, sgRNAs predicted to insert missense mutations were selected, and when multiple sgRNAs were predicted to target the same residue, the highest LFC value was considered. Non-targeted residues were presented in grey.

### Cell line genotyping

Knockout and knock-in efficiencies were evaluated using targeted amplicon sequencing. Briefly, cells transduced with the sgRNAs of interest were collected at day 4 post-antibiotic selection. Genomic DNA was extracted with QuickExtract (LGC Biosearch Technologies) according to the manufacturer’s instructions. Target genomic loci were PCR-amplified using Q5 high-fidelity DNA polymerase (New England Biolabs) and primer pairs designed with Primer3Plus (**Table S5**) (37). Genomic DNA from *AAVS1*-targeted cells was used as a WT reference sequence. Samples were sequenced at Genome Québec. For base editing experiments, sequencing traces were analyzed for in-window and out-of-window single-base substitutions using the Inference of CRISPR Editing (ICE) analysis tool (v3.0) (EditCo Bio). For CRISPR-KO experiments, editing efficiencies were computed using TIDE (38).

### Cell Viability Assays

HCT116 or RKO cells were seeded in duplicate at 500 cells/well (HCT116) or 1,500 cells/well (RKO) in black/clear-bottom 96-well plates (Corning) in 100 µL complete DMEM. After overnight incubation, threefold serial dilutions of HRO761 or VVD-214 were applied to the wells to achieve final concentrations ranging from 0.001 to 30 µM. After a 7-day incubation period, cell confluency was measured through two methods. When available, confluency values were computed via direct imaging on an Incucyte system (Sartorius AG). Alternatively, cells were fixed and permeabilized in 2% (w/v) PFA/0.5% (v/v) Triton X-100 in PBS for 10 minutes, washed three times with PBS–0.1% (v/v) Tween 20 (PBST), then stained with 1µg/mL 4, 6-diamidino-2-phenylindole (DAPI) for 5 minutes and washed twice with PBST. Plates were imaged on a Cytation C10 on a high-resolution fluorescence configuration with a 10X objective (Agilent Technologies), acquiring 9 images per well. Plates were analyzed with CellProfiler (39) to determine total DAPI intensities, reflective of well confluency, and data were normalized to DMSO-treated conditions. Data were plotted in GraphPad Prism 10 (GraphPad Software, LLC) and fitted to a curve using an [Inhibitor] vs. response – Variable slope (four parameters) to compute IC50 values.

For drug synergy experiments, HRO761 was used at a range of 0 to 1 µM, GSK-2830371 at a range of 0 to 5 µM, and AZD-7648 at a range of 0 to 10 µM using a D300e digital dispenser (Tecan). Confluency values were computed via direct imaging on an Incucyte system (Sartorius AG), and data were normalized to DMSO-treated controls. Synergy scores were calculated using SynergyFinder 3.0 (40), using the ZIP method considering outlier detection, four parameters (LL4) curve fitting, and ZIP correction.

### Clonogenic survival assays

Cells were seeded in 6-well plates at the following densities (in cells/well):

MSI-high: HCT116, 500; SW48, 1000; LoVo, 2000; SNU638, 2000; RKO, 500; LS411N, 2500. MSS/MSI-low: HEC1B, 750; NCIH747, 1000; JEG3, 1500; JAR, 1000; ESS1, 1750; NUGC3, 1000.

Twenty-four h later, cells were exposed to HRO761 at 100 nM. Cultures were maintained until distinct colonies formed, defined as approximately >50 cells per colony. Colonies were rinsed with PBS and stained with 0.4% (w/v) crystal violet in 20% (v/v) methanol for 30 min. The stain was aspirated, plates were rinsed twice with double-distilled H_2_O, air-dried, and colonies were quantified using a GelCount instrument (Oxford Optronix). Experiments were performed in technical duplicates. Data was represented as the surviving fraction of colonies when compared to the DMSO-treated control.

### Immunoblotting

Cells were lysed in Benzonase buffer [25 mM Tris (pH 8.8), 40 mM NaCl, 0.05% (w/v) SDS, 2 mM MgCl_2_, 10 U/mL Benzonase (Millipore Sigma)] or in M-PER extraction buffer (Life Technologies) supplemented with Pierce protease inhibitor tablets (Thermo Fisher Scientific) for 90 min. Samples were clarified for 15-20 min at 13,000g, and supernatants were diluted in NuPAGE™ LDS Sample Buffer (4×) and boiled at 95 °C for 5 min (Life Technologies). Equivalent protein amounts were subjected to gel electrophoresis, and proteins were transferred onto 0.45 µm nitrocellulose membranes using the Mini-PROTEAN electrophoresis and wet transfer system (Bio-Rad). Membranes were incubated overnight at 4 °C in primary antibodies in 3% (w/v) BSA–PBST. Primary antibodies used include WRN (Cell Signaling, 4666, 1:1,000), pKAP1 (Ser824, Cell Signaling, 4127, 1:1,000), KAP1 (Cell Signaling, 4124, 1:1,000), pATM (Ser1981, Cell Signaling, 13050, 1:1,000), ATM (Cell Signaling, 2873, 1:1,000), pCHK2 (Thr68, Cell Signaling, 2661 or 2197, 1:1,000), CHK2 (Millipore Sigma 05-649 or Cell Signaling 3440, 1:1,000), Tubulin (Cell Signaling, 2148, 1:2000) and β-actin (Invitrogen, MA1-140, 1:5,000). Proteins were detected using HRP-conjugated anti-mouse and anti-rabbit IgG secondary antibodies (1:5,000, Jackson Laboratories) and enhanced chemiluminescence (SuperSignal West Pico PLUS, Life Technologies). Signals were acquired on a ChemiDoc system (Bio-Rad).

### Deep mutational scanning

We performed deep mutational scanning of WRN (UniProt: Q14191) by creating a barcoded single-amino acid substitution library covering exons 19 (amino acids 697-758), 21-23 (amino acids 817-942) of the full-length cDNA cloned in the pLV[Exp]-Puro-CMV at VectorBuilder. Barcode–variant mapping was derived from plasmid pre-sequencing (**Table S2**). Target HCT116 cells were transduced at an MOI of ~0.3 to favor single integrations. The screen was conducted in technical triplicates, and library coverage of >400 cells per gene variant was maintained at every step. Puromycin-containing medium (2 µg/mL) was added 2 days after infection to select for transductants. Selection was continued until 96 h after infection, which was considered the initial time point (T0), after which cells were exposed to vehicle (DMSO), HRO761 (40 nM), and VVD-214 (7 nM) every 4 days for a total of 12 days (T12). Genomic DNA was isolated from endpoint (T12) cell pellets using the QIAamp Blood Maxi Kit (Qiagen), and barcode sequences were amplified by PCR using NEBNext Ultra II Q5 Master Mix (New England Biolabs). The i5 and i7 multiplexing barcodes were added in a second round of PCR, and final gel-purified products were sequenced on an Illumina NextSeq500 system at the Lunenfeld-Tanenbaum Research Institute (LTRI-NBCC).

Data were analyzed using R version 4.5.1. To determine the representation of WRN variants, barcodes were first aligned to the library reference sequences using Bowtie (41) to generate read counts for each sample replicate and library member. LFCs were then calculated by comparing WRN inhibitor-treated vs untreated populations at the endpoint (T12) using DESeq2 (1.48.2; 42). LFCs and p-values were calculated from normalized counts per library depth using a model that included replicate batch effects to mitigate potential bias. Weighted effect scores per amino acid change were computed by first multiplying LFCs by the negative log10 of the adjusted p values. Then, the mean effect scores per-residue was then standardized (z-score calculation), such that the reported value represents the number of standard deviations each value lies from the overall mean ((x − μ)/σ).

Data were represented as heatmaps or correlation plots using ComplexHeatmap 2.24.1 (43). Per-residue mean weighted effect scores (standardized) were mapped onto the corresponding WRN structures bound to HRO761 (PDB ID: 8PFO, 9), or VVD-214 (PDB ID: 7GQU, 8) using UCSF ChimeraX version 1.10rc202506130232 (36). Non-targeted amino acids were represented in gray.

### Immunofluorescence

Cells were seeded at a density of 2 x 10^4^ cells per well of black clear-bottom 96-well plates. After overnight culture, cells were treated with HRO761 (10 μM), VVD-214 (1 μM) or left untreated for 24 h. Cells were then fixed and permeabilized for 10 min using a solution of 2% (w/v) paraformaldehyde and 0.5% (v/v) Triton X-100 in PBS. After blocking with TBS–0.1% (v/v) Tween 20 (TBST) containing 3% (w/v) BSA for 1 h, incubated with a primary antibody against H2AX phospho-Ser139 (γH2A.X) (1:5,000, Biolegend, 613402) for 1 h at room temperature. Cells were washed three times with PBS and incubation with secondary antibody anti-mouse Alexa Fluor 594 (Life Technologies) was then performed for 45 min. Cells were stained with 1 μg/mL DAPI for 10 min for nuclear staining and imaged using a BioTek Cytation C10 imaging reader with a 40× objective (Agilent Technologies). Quantification of γH2A.X integrated nuclear intensity was performed using CellProfiler (39). All experiments were performed in biological triplicate, and data were represented as SuperPlots using GraphPad Prism 10 (GraphPad Software, LLC).

### PRISM drug-response and Cancer Dependency Map analyses

HRO761 was submitted to the Broad Institute for drug–response analyses as part of the PRISM project (**Table S3**). Data from the Cancer Dependency Map, including WRN Chronos scores (DepMap Public 25Q3) and microsatellite instability status from the Cancer Cell Line Encyclopedia (CCLE), were downloaded from the DepMap website (**Table S3**). Cell lines were matched based on their official names, and only instances with complete PRISM and WRN dependency data were preserved (N = 678 cell lines). To represent the relationship between HRO761 sensitivity and WRN dependency, PRISM AUC values were used. When data per cell line were duplicated, the average AUC was calculated. Data were plotted using GraphPad Prism 10 (GraphPad Software, LLC).

### CRISPR/Cas9 chemogenomic screening

CRISPR/Cas9 screens were performed as previously described (33), with slight modifications. Cells were transduced with lentivirus carrying the TKOv3 sgRNA library at a multiplicity-of-infection (MOI) of ~0.3. The screen was conducted in technical duplicates, and library coverage of >200 cells per sgRNA was maintained at every step. Puromycin-containing medium (1.0 µg/mL) was added 2 days after infection to select for transductants. Selection was continued until 96 h after infection, which was considered the initial time point (T0). HRO761 was added to the cells starting from day 6 (T6) at 100 nM, corresponding to ~LD20. From T10 onwards, HRO761-containing medium was subsequently refreshed every four days until the screen was terminated at T18. To identify genes whose deletion sensitizes cells to HRO761, genomic DNA was isolated from surviving cells using the QIAamp Blood Maxi Kit (Qiagen), and genome-integrated sgRNA sequences were amplified by PCR using NEBNext Ultra II Q5 Master Mix (New England Biolabs). i5 and i7 multiplexing barcodes were added in a second round of PCR, and final gel-purified products were sequenced on an Illumina NextSeq500 system to determine sgRNA representation in each sample. Read counts per sgRNA were obtained using the MAGeCK count command. Normalized Z scores (NormZ) were obtained by comparing DMSO-treated vs HRO761-treated cells at the end point (T18) using DrugZ. Data was represented as a gene ranking dot plot with ggplot2 version 4.0.0.

## Declarations

### Availability of data and materials

NGS data for base editing, DMS and chemogenomic screens are deposited in the NCBI Sequence Read Archive (SRA) under BioProject ID: PRJNA140371. Raw data for all experiments will be deposited in Figshare upon publication. Cell lines and plasmids are available upon request.

### Authors’ contributions

MEO performed base editing and DMS screens, analyzed G729S/+ lines and WRN overexpressing mutants, and performed the chemogenomic screen validation. NL performed the chemogenomic screens and drug synergy experiments. WM analyzed the differential expression of WRN overexpressing mutants *via* survival analyses and immunofluorescence. SS generated and analyzed the F730S/+ cells. VC-T, JB, and CF analyzed and represented data. AM-OP, J-JC-R, and TS provided experimental support and reagent generation. JTFY provided scientific guidance and feedback on the study. AA-Q conceptualized the study, supervised MO and NL, and co-wrote the manuscript. RC-M supervised MO, WM, VC-T, AM-OP, J-JC-R, and TS, analyzed and represented data, and co-wrote the manuscript. All authors edited and approved the manuscript.

## Acknowledgements

The authors thank Ms. Tania Schramek of the Victor Phillip Dahdaleh Institute of Genomic Medicine for her contributions to the clarity and refinement of the manuscript. The authors thank Luke Dow and Francisco Sanchez-Rivera for advice on screen implementation. The authors thank Deniz Meneksedag-Erol, Golshad Ghojoghi, and Felipe Veloso for structural insights, and all members of the Cuella Martin laboratory for support and helpful discussions.

## Supplementary material

Supplementary Figures S1-S3

Table S1. Raw read counts and LFCs for base editing screens in HCT116 cells with the sgRNA annotation and information on designed negative and positive controls.

Table S2. Read counts and LFCs for DMS screens in HCT116 cells with construct annotation.

Table S3. PRISM dataset with AUC values for HRO761 across cell lines and DepMap WRN dependency and MSI status (2025Q3).

Table S4. Readcounts and NormZ values for the chemogenomic screen in RKO cells.

Table S5. Oligonucleotide sequences, including sgRNA sequences and primer pairs used for editing efficiency determination.

