## Supplementary Material for "Anticipating on-target resistance to WRN inhibitors in microsatellite unstable cancers"

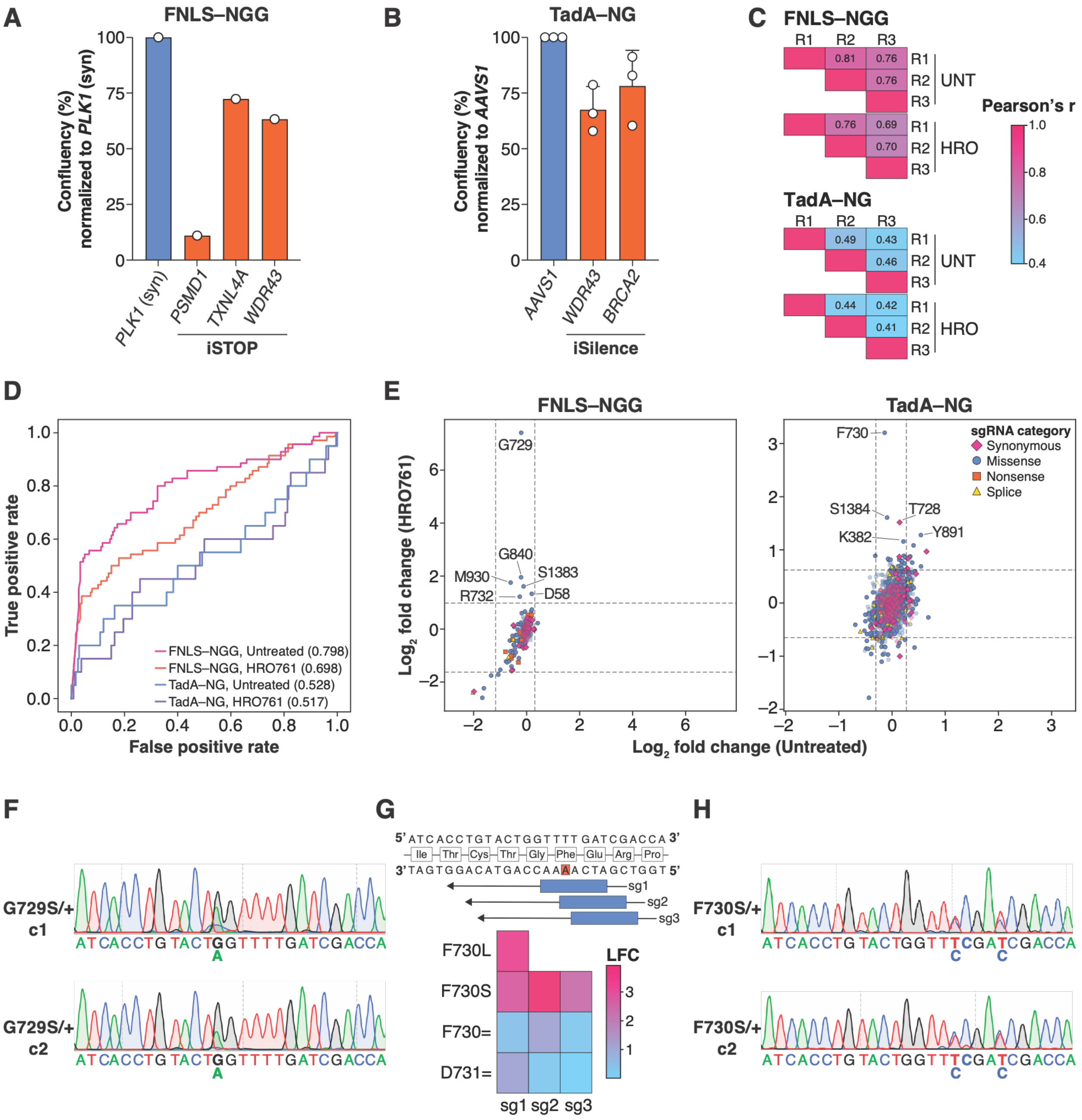


**Figure S1**. (**A**) HCT116–FNLS–NGG cells were transfected with synthetic sgRNAs inserting stop codons (iSTOP) in *PSMD1*, *TXNL4A,* and *WDR43*. An sgRNA introducing a synonymous mutation on *PLK1* was used as a control. Confluency was measured using Incucyte. N = 1. (**B**) HCT116–TadA–NG cells were transfected with synthetic sgRNAs targeting *AAVS1* or with sgRNAs targeting the start codons (iSilence) of *WDR43* or *BRCA2*. Confluency was measured using Incucyte. Mean ± SD. N = 3. (**C**) Heatmap of Pearson (r) correlation values for FNLS–NGG and TadA–NG screens. (**D**) ROC-AUC analyses of screen performance. True positives include iSTOP and iSilence sgRNAs targeting essential genes for FNLS–NGG and TadA–NG, respectively. True negatives include empty-window (no edits) and designed negative controls (**Table S1**). **(E)** Correlation plot of LFCs in untreated *vs* HRO761-treated base editing screens. Dotted lines represent thresholds for biological relevance. (**F**) Sanger sequencing traces for the G729S/+ clones used in Fig. 1F, G. (**G**) Evaluation of mutational outcomes induced by sgRNAs targeting the F730 codon in the presence of HRO761. Edited HCT116–ABE8e–SpG cells were analyzed by Sanger sequencing and then treated with 600 nM HRO761 for 8 days or left untreated. Surviving cells were Sanger sequenced, and traces were analyzed using the ICE analysis tool (v3.0) (EditCo Bio). LFC values were calculated by comparing the percentage of edited alleles in treated versus untreated conditions. (**H**) Sanger sequencing traces for the F730S/+ clones used in Fig. 1H, I.

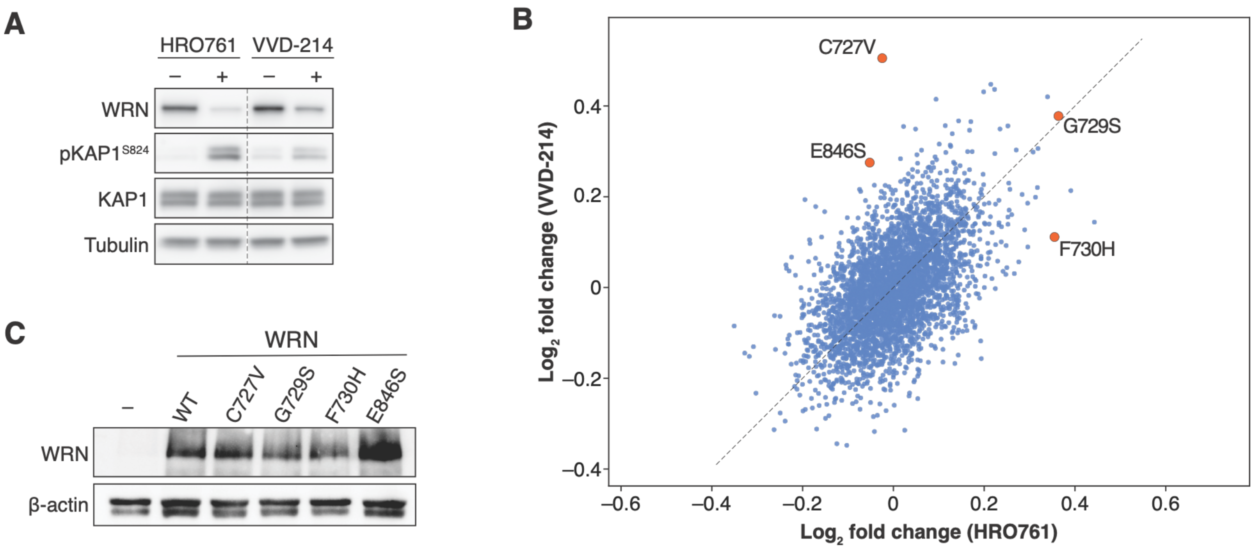


**Figure S2.** (**A**) Immunoblot of the inhibitor-induced degradation of WRN and the activation of DNA damage in HCT116 cells treated with HRO761 (1 µM) and VVD-214 (1 µM) for 24 h. (**B**) Correlation plot of LFC values for the HRO761 and VVD-214 DMS experiments. Mutations selected for validation are highlighted. (**C**) Immunoblot showing expression of WRN mutants in HCT116 cells used for the experiments in Fig. 3.

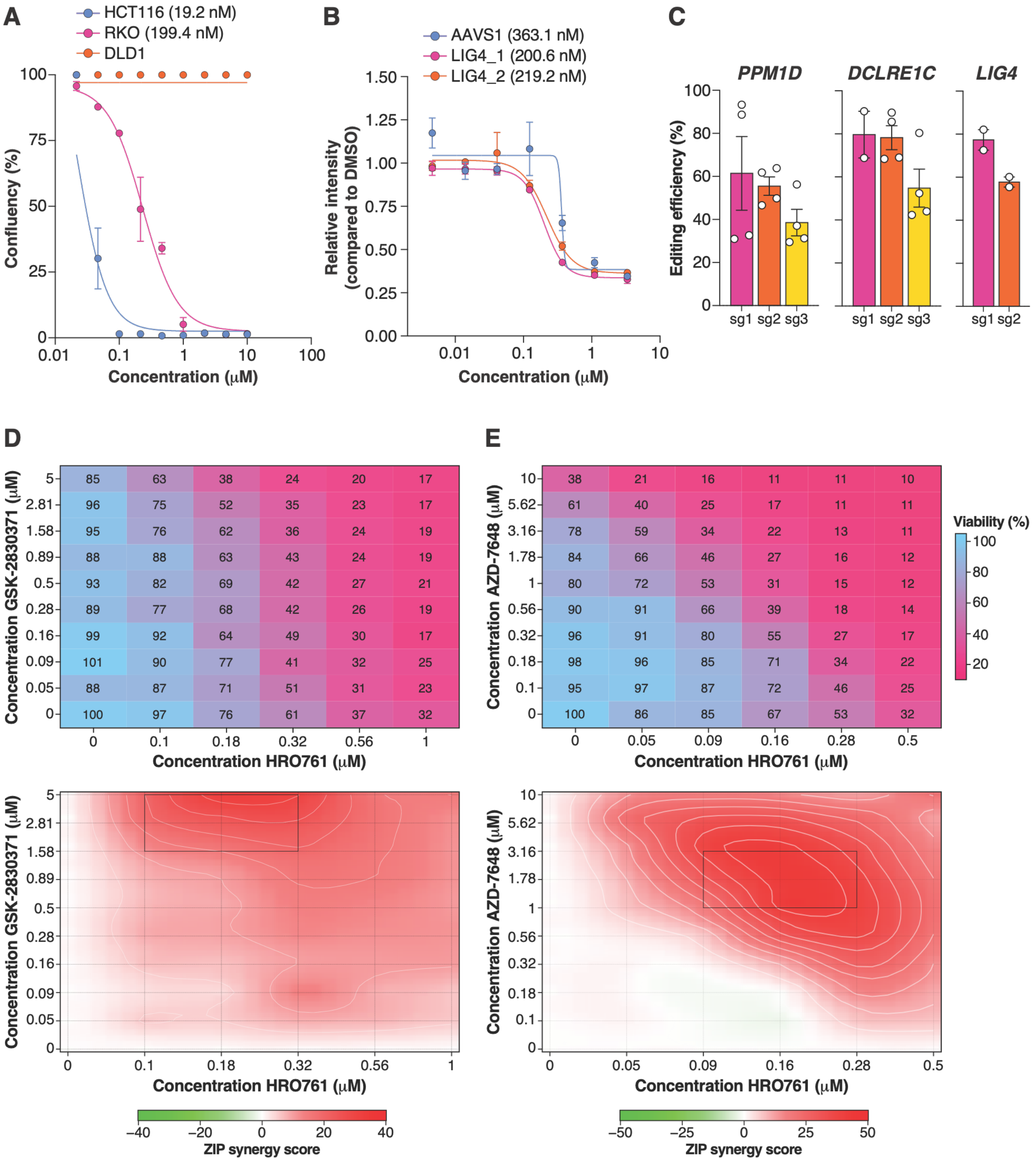


**Figure S3.** (**A**) Survival curves of HCT116, RKO and DLD1 cells treated with HRO761. Cells were exposed to HRO761 for 12 days, and confluency was measured using Incucyte. Mean ± s.e.m. N = 2. (**B**) Survival curves of RKO cells transduced with sgRNAs targeting *LIG4* using a one-vector system. Six days post-infection, cells were exposed to HRO761 for 7 days, and confluency was measured by total DAPI intensity. Mean ± s.e.m. compared to DMSO-treated control cells. N = 2. (**C**) Editing efficiencies for sgRNAs used in the survival assays from **4E** and **S3B.** Mean ± s.e.m. N = 2-4. (**D, E)** Combinatorial treatment matrix of RKO cells treated with HRO761, GSK-2830371 (D), or AZD-768 (E) for 7 days. Confluency was measured with Incucyte. Relative survival is calculated with respect to an untreated control. Mean for N = 2. Corresponding ZIP synergy scores are depicted below, with areas of maximum synergy highlighted.

### Supplementary Tables

**Table S1**. Raw read counts and LFCs for base editing screens in HCT116 cells with the sgRNA annotation and information on designed negative and positive controls.

**Table S2**. Read counts and LFCs for DMS screens in HCT116 cells with construct annotation.

**Table S3**. PRISM dataset with AUC values for HRO761 across cell lines and DepMap WRN dependency and MSI status (2025Q3).

**Table S4**. Readcounts and NormZ values for the chemogenomic screen in RKO cells.

**Table S5.** Oligonucleotide sequences, including sgRNA sequences and primer pairs used for editing efficiency determination.
